# Locomotor effects of a fibrosis-based immune response in stickleback fish

**DOI:** 10.1101/2023.06.24.546342

**Authors:** David G. Matthews, Meghan F. Maciejewski, Greta A. Wong, George V. Lauder, Daniel I. Bolnick

**Affiliations:** Organismic and Evolutionary Biology, Harvard University, Cambridge, 02138, MA, USA; Museum of Comparative Zoology, Harvard University, Cambridge, 02138, MA, USA; Department of Evolution, Ecology, and Behavior, University of Illinois at Urbana-Champaign, Champaign, 61820, IL, USA; Department of Ecology Evolutionary Biology, University of Connecticut, Storrs, 06269, CT, USA

**Keywords:** Stickleback, Immune response, C-start, performance, Structural equation modeling

## Abstract

The vertebrate immune system provides an impressively effective defense against parasites and pathogens. However, these benefits must be balanced against a range of costly side-effects including energy loss and risks of auto-immunity. These costs might include biomechanical impairment of movement, but little is known about the intersection between immunity and biomechanics. Here, we show that a fibrosis immune response in threespine stickleback (Gasterosteus aculeatus) has collateral effects on their locomotion. When freshwater stickleback are infected with the tapeworm parasite Schistocephalus solidus, they face an array of fitness consequences ranging from impaired body condition and fertility to an increased risk of mortality. To fight the infection, some stickleback will initiate a fibrosis immune response in which they produce excess collagenous tissue in their coelom. Although fibrosis is effective at reducing infection, some populations of stickleback actively suppress this immune response, possibly because the costs of fibrosis outweigh the benefits. Here we quantify the locomotor effects of the fibrosis immune response in the absence of parasites to investigate whether there are collateral costs of fibrosis that could help explain why some fish forego this effective defense. To do this, we induce fibrosis in stickleback and then test their C-start escape performance. Additionally, we measure the severity of fibrosis, body stiffness, and body curvature during the escape response. We were able to estimate performance costs of fibrosis by including these variables as intermediates in a structural equation model. This model reveals that among control fish without fibrosis, there is a performance cost associated with increased body stiffness. However, fish with fibrosis did not experience this cost but rather displayed increased performance with higher fibrosis severity. This result demonstrates that the adaptive landscape of immune responses can be complex with the potential for wide reaching and unexpected fitness consequences.

## Introduction

When animals contract parasites, they often exhibit an immune response in an attempt to minimize the deleterious effects or stave off the parasite altogether. One example of a well-studied host-parasite system is the case of Schistocephalus solidus, a tapeworm parasite, infecting freshwater populations of threespine stickleback fish (Gasterosteus aculeatus). When these cestode parasites severely infect a fish, they have been found to impose many costs including impaired host body condition (Tierney et al. 1996; Barber and Svensson 2003; Jolles et al. 2020), reduced fertility (Schultz et al. 2006), reduced steady swimming speed and acceleration (Blake et al. 2006; Jolles et al. 2020), reduced ability to school (Jolles et al. 2020), and reduced behavioral antipredator responses (Giles 1983; Milinski 1985). Some of these costs are attributable to the energy lost to the parasites and others are due to behavioral shifts that the parasite induces to help it complete its life cycle by facilitating bird predation (Grécias et al. 2016, 2018, 2020; Berger and Aubin-Horth 2020). Stickleback are able to mount an immune response to help fend off the parasites, and one aspect of this response is peritoneal fibrosis, a dense fibrous layer of collagenous tissue laid down inside the coelom to physically contain the parasite (Lohman et al. 2017; Weber et al. 2017, 2022; Vrtílek and Bolnick 2021). This response is also strongly associated with the formation of granuloma cysts that encase and often kill the cestode (Weber et al. 2017, 2022). Fibrosis is known to be an effective immune response, with fibrotic fish containing smaller and fewer tapeworms (De Lisle and Bolnick 2021; Weber et al. 2022). Additionally, natural populations with this immune response have a lower prevalence of the parasite (Hund et al. 2022; Weber et al. 2022). Surprisingly, some populations of stickleback forego this immune response, letting the parasite grow relatively unchecked (De Lisle et al. 2022; Weber et al. 2022). Many of these populations up-regulate fibrosis-suppression genes, indicating that they are actively suppressing this immune response (Lohman et al. 2017; Fuess et al. 2021). Furthermore, there is evidence of positive selection favoring fixation of large deletions within pro-fibrotic genes (Weber et al. 2022). This immune suppression suggests that there is a cost to the peritoneal fibrosis itself that could sometimes outweigh the cost of the parasite (De Lisle et al. 2022). Although this is an ongoing area of research, we know that the deleterious effects of fibrosis include lowered foraging success (De Lisle and Bolnick 2021) and a reduction of fertility in both males and females (De Lisle and Bolnick 2021; Weber et al. 2022). It is also likely that there is an energetic cost associated with creating fibrotic tissue. In addition to these costs, we would expect that the fibrous tissue would change the material properties of fish’s bodies, as fibrotic tissue has been found to change the stiffness and viscoelastic properties of many tissues across numerous mammalian models (Wells 2013; Zhu et al. 2016). Additionally, we know that body stiffness and non-uniform distribution of material properties both have a large impact on swimming performance (Long and Nipper 1996; Long et al. 1996; Long 1998; Lucas et al. 2015; Luo et al. 2020). Therefore, we would expect that the material property changes associated with fibrosis would impact the swimming ability of an affected fish. While stickleback primarily swim steadily by rowing their pectoral fins (Walker and Westneat 2002; Walker 2004), they are still susceptible to these effects as they rely on locomotor forces generated by the whole body during escape responses and turning behaviors when considerable body bending occurs. C-start escape responses are one locomotor behavior in stickleback that relies on bending of the body to generate propulsive forces. Fast-start behaviors such as the C-start are a well-known and widely studied behavior since they are critical for survival and are extremely widespread among fish (Eaton and Hackett 1984; Domenici and Blake 1997; Domenici and Hale 2019). The behavior consists of two main portions 1: Stage 1, when fish bend their body about the center of mass until their body is shaped like a “C”, causing them to reorient; and Stage 2, where fish unfurl their tail to propel themselves forward in their new orientation. Although this behavior was once thought to be highly stereotyped both interspecifically and intraspecifically (Eaton et al. 1977; Webb 1978), the escape behavior has proved to be more variable in both regards than previously recognized as more species have been examined with larger sample sizes (Marras et al. 2011; Domenici and Hale 2019). However, individual escape performance is highly repeatable (Marras et al. 2011), indicating that the variation in escape performance has some innate basis and can be acted on by selection.

Past robotic and simulation studies have identified body stiffness as an important variable for determining C-start performance, and one previous result is that intermediate stiffnesses leads to optimal escape performance (Ahlborn et al. 1997; Witt et al. 2015). Generally, bodies that are too flexible do not produce high enough forces, while bodies that are too stiff do not create the necessary thrust-producing vortices to power the escape. Within this intermediate range, the ideal stiffness value also depends on the stage of the escape response. Currier and Modarres-Sadeghi (2019) found that robotic models maximized total turn angle in Stage 1 when they had a more flexible body, but the same models maximized acceleration away from the stimulus in Stage 2 when the body was stiffer. Interestingly, fish have been shown to antagonistically contract their body musculature to modulate stiffness during swimming and escape responses (Long and Nipper 1996; Long 1998; Tytell and Lauder 2002; Tytell et al. 2018). This could allow them to actively tune their body stiffness between different portions of the escape response. Here we ask whether the fibrosis immune response in stickleback has any locomotor consequences that could impact the adaptive value of immunity against the Schistocephalus parasite. Specifically, we investigate the effects of artificially induced fibrosis, thereby removing the confounding effect of parasitic infection to help disentangle the selective pressures that have led to immune suppression in some populations. We focus on C-starts because this behavior involves a high degree of body bending and because it has high fitness consequences as a predator avoidance mechanism (Langerhans et al. 2004; Langerhans and Reznick 2010). We also know that there is phenotypic variation in C-start performance, as comparisons of freshwater vs. marine stickleback (Taylor and McPhail 1986) and between different freshwater populations (Andraso and Barron 1995) have shown divergence over small temporal and spatial scales that were attributed to local selective pressures. Additionally, C-start performance is important for the completion of this trophically transmitted parasite’s life cycle as the parasite breeds in the intestines birds that have eaten infected stickleback. Therefore, the host’s and parasite’s interests are in conflict as the host wants to evade predation and the parasites want to facilitate it. We predict that fish with higher levels of fibrosis will have decreased performance during Stage 1 of the C-start behavior (lower angular velocity) and increased performance during Stage 2 (higher linear velocity). Specifically, we hypothesize a multifaceted mechanism for this effect in which fibrosis increases body stiffness, stiffness decreases the degree of body curvature during the C-start, and low levels of body curvature will decrease their ability to reorient in Stage 1 while increasing linear velocity in Stage 2 (Supplemental Figure 1A).

## Methods

We tested our hypothesized mechanisms linking fibrosis to escape performance (Supplemental Figure 1) by first measuring each proposed variable. To do this we induced fibrosis without the presence of Schistocephalus parasites in an experimental group of fish (with a control group for comparison), recorded them performing C-start escape responses, analyzed the videos, measured their body stiffness, and then dissected them to obtain a fibrosis severity score (Figure 2). These data were then included in a structural equation model (SEM), a category of statistical models that includes path analysis, to understand the interrelationships between these variables, and ultimately to estimate the effect of fibrosis on escape performance.

**Fig. 1.**
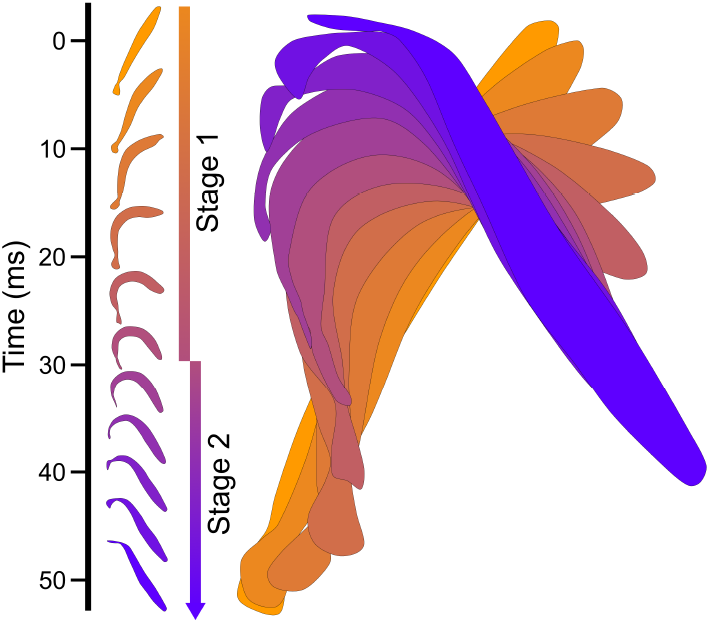
Example C-start escape response of a stickleback showing the body position through time. Time points are captured 5ms apart and time is represented as a change in color. Non-overlapping body outlines demonstrate the change in body bending over the course of the escape response without relative spatial context. Overlapping body outlines represent the true position of the fish in space over the course of the escape response.

**Fig. 2.**
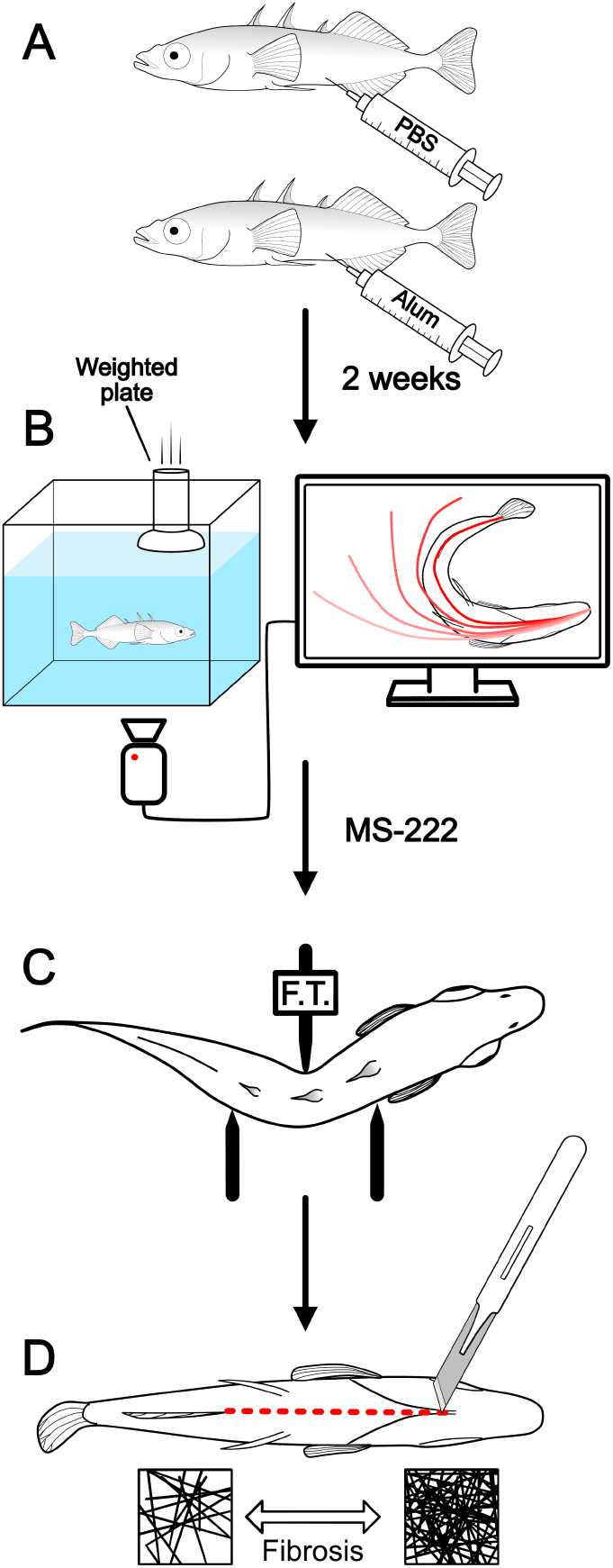
Overview of the methodology used in this study. A) We injected threespine stickleback (*Gasterosteus aculeatus*) with either phosphate buffered saline (PBS) as a control or with alum, an immune adjuvant that stimulates a fibrosis response. B) Two weeks later, we filmed the fish ventrally as a weighted plate was dropped in their tank to stimulate a C-start escape response. We recorded each sequence then traced the midline to analyze kinematics. C) After euthanizing the fish with MS-222, we measured their body stiffness using a three-point bending test and a force transducer (F.T.). D) We dissected the fish and evaluated the amount of fibrous tissue present in the body cavity.

### Specimen collection and Injections

We collected adult threespine stickleback (Gasterosteus aculeatus) from Roselle Lake on Vancouver Island, British Columbia in June of 2018. We stripped eggs from gravid females and fertilized these with macerated testes to generate embryos, which were transported to the University of Connecticut and reared to maturity. We randomly selected 20 adult lab-raised fish as the experimental population and 12 fish as the control population. We then induced fibrosis in the experimental population following Hund et al (2022). In short, we injected 20 µL of an immune adjuvant, a 1% solution of alum in PBS, into the peritoneal cavity of the experimental fish. Alum is used to induce fibrosis without the presence of parasites (Figure 2A) as it induces an inflammatory response that leads to peritoneal fibrosis, exactly mimicking the fibrosis seen in cestode-infected fish (Vrtílek and Bolnick 2021; Hund et al. 2022). Importantly, this allowed us to measure the effects of fibrosis without confounding effects from the parasites such as increased energy demands and behavioral manipulation of the host by secretory-excretory products. Simultaneously, we injected the control fish with an equivalent volume of 1X phosphate buffered saline (PBS) to account for any effects of the injection itself. After waiting 10 days we repeated the same injections on the contralateral side to maximize the immune response. We then returned the fish to their tanks for two weeks as this is sufficient time for fibrosis to fully develop, but not enough time that fish begin breaking down the fibrotic tissue (Hund et al. 2022).

### Data collection

At the end of the two weeks fish were isolated in individual filming tanks. One at a time they were placed in a dark room in front of two high speed video cameras (Photron AX50, 1024 x 1024 pixels, Photron Inc.), one recording a lateral view and the other recording a ventral view. After allowing the fish to acclimate, we illuminated the tank with infrared lights and induced C-start escape responses by dropping a weighted plate directly in front of the fish (Figure 2B, Supplemental video 1). We attached the weighted plate to a monofilament line so that it would stop shortly after contacting the surface of the water, thereby creating a pressure wave and startling the fish, but not creating substantial water flow within the tank. Each escape response was recorded at 1000 frames per second (1/2000 s shutter speed, 30mm focal length, f/4), and only analyzed if the side view revealed that the fish was parallel to the ground during the behavior. After each escape response the fish was left undisturbed until it returned to a calm state, indicated by decreased opercular motion and slow steady swimming within the tank. At this point, a new escape response was elicited and recorded. This process was repeated with each fish until their reaction to the weighted plate stimulus was diminished, as determined by elongated reaction times and reduced turning angle during the escape response. We did not use any videos in which the response was visibly diminished. Each set of videos was calibrated with a ruler held in the plane of focus of both cameras. After recording each fish, we immediately euthanized them in a buffered MS-222 solution. Fish were left in the solution for 20 minutes after the cessation of opercular motion, at which point we measured their passive body stiffness (Figure 2C). First, we clipped their pelvic spines flush with the body surface so that the external spines would not affect our measurement of body stiffness. Then we placed the fish in a submerged three-point bending apparatus with the middle arm attached to a force transducer (FUTEK LSB200; FUTEK Advanced Sensor Technology Inc., Irvine, CA USA). The two outer arms were adjustable and were positioned such that the first point was aligned with the base of the first dorsal spine and the second point was aligned with the anterior-most fin ray of the second dorsal fin (Supplemental Figure 2C). We positioned the middle arm at the mid-point of the other two arms and lowered it until it contacted the stickleback such that the fish’s midline was straight. We then began continuous force recording (at 1000 Hz), measuring the force at the starting point. Using a micrometer, we lowered the middle arm in 0.5mm increments from 0 to 3.5mm, allowing the force measurement to stabilize at each position. Additionally, at each position we photographed the fish from a dorsal view so that we could measure body curvature. Finally, we removed the fish from the bending apparatus and photographed them head on (Supplemental Figure 2D) so that their cross-sectional area could be measured. Each fish was then immediately frozen at -20°C. We later thawed the fish and dissected them to measure the severity of their fibrosis response (Figure 2D; measured on a scale from 0 to 4, following Hund et al. 2022; See Supplemental Video 1 from Hund et al. 2022).

### Data analysis

We first used high-speed videos of C-start responses to measure body curvature and escape performance during both Stage 1 and Stage 2 of the escape response. Since we were interested in the maximal body curvature during an escape, we began by estimating the frame in which maximum bending was achieved and then sampled 5 frames before and after. Within each of these frames we used a custom MATLAB script to digitize the midline from the snout to the caudal peduncle. We then found the coordinates of the points on the midline that were 1/3, 1/2, and 2/3 of the distance from snout to peduncle (Supplemental Figure 2A). These points roughly correspond to the position of the coelom, so we expect any effect on curvature due to variation in fibrosis and stiffness to be most pronounced here. We then used these three points to calculate curvature, where curvature is the inverse of the radius. We measured curvature in all the frames and used the maximum value among them as the peak curvature achieved during that particular escape response. After collecting these data for all recorded C-starts, we corrected for body size by regressing curvature on standard length of each fish and recording the residuals. Next, we measured the escape performance during each individual C-start. The escape response is divided into two stages: Stage 1 when fish bend their body to change orientation, and Stage 2 when they straighten their body again and accelerate away from the stimulus (Figure 1). First, we identified three frames in each video corresponding to (1) the frame immediately preceding rotation of the head at the beginning of Stage 1; (2) the frame immediately preceding extension of the tail, marking the transition from Stage 1 to Stage 2; (3) the frame 8ms (eight frames) after the transition to Stage 2. We chose to only use the first 8ms of Stage 2 to minimize behavioral variation occurring at later stages of C-starts (Domenici and Hale 2019). In each of these three frames we then digitized 3 landmarks along the midline corresponding to the anterior tip of the head, the posterior-most point where left and right branchiostegal rays overlap, and the point along the midline nearest to the pectoral fins (Supplemental Figure 2B). The third landmark is a good approximation of the center of mass in stickleback (Andraso and Barron 1995). We measured the fish’s heading at each time point by finding the line between the first and second landmarks. We defined Stage 1 performance as the average angular velocity over Stage 1, measured as the change in heading between the first and second subsampled frames divided by the time elapsed between the two frames. We measured Stage 2 performance by finding the distance between the third landmark (center of mass) in the second subsampled frame and the same landmark in the third frame, then dividing this distance by eight milliseconds. This gave the average linear velocity at the beginning of Stage 2 of the C-start response. Both the angular velocity in Stage 1 (Walker et al. 2005) and the linear velocity in Stage 2 (Katzir and Camhi 1993; Walker et al. 2005) have been found to directly impact likelihood of a successful escape and are therefore relevant performance metrics. To convert the force data from the three-point bending test into a stiffness value we first measured the curvature of each fish’s body at each displacement (0-3.5mm). To accomplish this, we digitized three landmarks in each picture, corresponding to the points along the midline where the three arms of the bending apparatus contacted the fish (Supplemental Figure 2C). These points are in similar positions to those used to measure curvature in escape response videos, meaning that stiffness and body curvature are measured over the same body span. We then calculated the curvature of the body by finding the curvature of the three landmarks. Next, we used our force recordings to find the maximum force observed at each displacement value of the middle point (0-3.5mm). The force observed at zero displacement was then subtracted from each value to account for buoyancy of the fish in the testing apparatus. To linearize the data, we log-transformed bending forces and regressed these against the measured body curvature for each fish. We then found the slope of this semi-log plot as a measurement of stiffness, similar to Young’s modulus. Finally, we size corrected these values by regressing stiffness on the cross-sectional area of each fish and recording the residuals. Cross sectional area was measured in Image-J (V1.53e) by fitting ovals to the body outline in anterior photos of each fish (Supplemental Figure 2D).

### Statistics

After visually examining the data, we saw that three experimental fish and one control fish had stiffness values that were well outside the range seen across all other fish, with two experimental fish being unusually flexible and the other two fish being unusually stiff. We used a Rosner test of multiple outliers (EnvStats V2.4.0 in R V4.0.2) (Millard 2013) to determine whether these four values were significantly outside the stiffness distribution measured from other individuals. All four points were deemed to be significant outliers (Supplemental Table 3), and because the response to alum treatment is sometimes incomplete (Hund et al. 2022) they were excluded from all further analyses. Additionally, we recalculated all size-corrected residuals with these four points removed. Finally, we linearly transformed fibrosis severity scores to lie on a scale from 0-1 so that coefficient estimates represented the effect of maximum fibrosis. We first confirmed that our alum treatment successfully induced a higher degree of fibrosis than the control injections by conducting a Wilcoxon signed rank test. This non-parametric approach accounts for the non-continuous nature of fibrosis severity scores. However, since SEM requires parametric modelling, we also confirmed that we would obtain similar results using a parametric approach, regressing fibrosis severity score on treatment (PBS vs. alum). We similarly test whether fibrosis severity score can be used in parametric models as an independent variable by both regressing stiffness on fibrosis severity scores and analyzing the same two variables with a Spearman’s rank correlation. In both cases we obtained similar results between the parametric and non-parametric approaches, so all further analyses are conducted using linear regression. Next, we tested whether escape performance was impacted by acclimatization to the stimulus by regressing both performance variables against trial number and fish ID. This model allows each fish to have a different mean performance value and asks whether there is a general downward trend as we carry out more trials. We did not find any evidence of acclimatization in escape performance, so we conclude that our kinematic data represent the range of performances that each individual is capable of. Since we are interested in how fibrosis constrains maximal performance, we continue our analysis using only the highest recorded value of each performance metric for each individual stickleback. We assume that this value represents the highest performance that an individual is capable of because maximal escape performance is highly repeatable within an individual (Fuiman and Cowan 2003; Marras et al. 2011; Hitchcock et al. 2015; Jornod and Roche 2015), meaning that the highest measured performance likely reflects their actual peak ability (Domenici and Hale 2019). We then regressed maximum performance from both Stage 1 and Stage 2 on treatment to see if treatment had a directly measurable effect on performance. To further improve the predictive power of this model, we ran a multivariate regression using fibrosis severity score, body stiffness, and body curvature to independently predict both performance measurements. However, none of these approaches yielded a significant effect of our treatment on performance, so we continued our analysis using SEM.

The first step in an SEM analysis is to create hypothesized paths that reflect the hierarchical relationship between all the variables in the analysis. We start with a base path model wherein fibrosis affects performance through stiffness and body curvature (Supplemental Figure 1A). After plotting all the individual regressions within this model, we also included an interaction between treatment and stiffness to better represent patterns of stiffness variation. We then created additional path models to reflect additional mechanisms through which fibrosis could affect performance. We propose two possible additional effects: a behavioral effect in which the body curvature is directly affected by fibrosis (Supplemental Figure 1B) and a force transfer effect where stiffness directly affects the fish’s ability to generate turning and propulsive forces (Supplemental Figure 1C). Finally, we created a model that included both the behavioral and the force transfer effects (Supplemental Figure 1D). We then ran all four path models using the “sem” function in the Lavaan package (V0.6-12, R V4.0.2) (Rosseel 2012). We compared all the models using a variety of fit indices (see Table 1) as well as the Akaike’s information criterion (AIC) and Bayesian information criterion (BIC). The fit indices were evaluated using the criteria set forth by Kline (2005). Since all three methods of evaluating best-fit indicated that the base model with a behavioral effect was the preferred model we obtained our final coefficient estimates from this model, including indirect and total effects. Specifically, we estimated coefficients for each line in the path diagram, each independent path that led from fibrosis to both performance metrics (indirect effects), and the summation of all paths connecting fibrosis to each performance metric (total effects). Additionally, we plotted the correlation represented by each line on the path diagram. Since fibrosis severity score is used to predict both stiffness and body curvature, we plot fibrosis severity score residuals from the body curvature regression against stiffness to account for the variance in fibrosis already explained by body curvature. Finally, we estimated effect sizes by dividing the total effect coefficient for each performance metric by the mean performance value of all individuals that lacked fibrosis. Since the fibrosis severity scores were scaled from zero to one this tells us the predicted percentage change in performance of going from no fibrosis to maximum fibrosis.

**Table 1.** Fit indices for the four tested SEM paths, including the degrees of freedom (DoF), the root mean square error of approximation (RMSEA), comparative fit index (CFI), goodness of fit index (GFI), and standardized root mean square residual (SRMR).

| Model | DoF | $\chi^2$ P-value<br>(>0.05) | RMSEA<br>(<0.08) | CFI<br>(>0.9) | GFI<br>(>0.95) | SRMR<br>(<0.08) | AIC | ΔAIC | AIC Weight | BIC | ΔBIC | BIC Weight |
| --- | --- | --- | --- | --- | --- | --- | --- | --- | --- | --- | --- | --- |
| Base model | 12 | 0.045 | 0.208 | 0.884 | 0.985 | 0.18 | -7.8 | 6.6 | 0.03 | 5.5 | 5.7 | 0.05 |
| + Behavior | 11 | 0.306 | 0.095 | 0.978 | 0.99 | 0.064 | -14.4 | 0 | 0.8 | -0.1 | 0 | 0.87 |
| + Force transfer | 10 | 0.025 | 0.241 | 0.87 | 0.985 | 0.17 | -4.7 | 9.7 | 0.01 | 10.4 | 10.6 | 0 |
| +Behavior<br>+ Force transfer | 9 | 0.218 | 0.134 | 0.964 | 0.99 | 0.057 | -11.3 | 3.1 | 0.17 | 4.8 | 4.9 | 0.08 |

## Results

Injecting stickleback with alum significantly increased fibrosis severity (mean fibrosis score from 0-1: PBS = 0.03, alum = 0.78) in both the parametric (p < 0.001) and non-parametric (p < 0.001) tests compared to control injections. Similarly, fibrosis severity significantly predicted body stiffness in both the parametric (p = 0.020) and non-parametric (p = 0.024) tests. These tests together confirm that we can include fibrosis severity score as a variable in all further parametric statistical tests. Furthermore, we found that there was no significant effect of trial number on maximum body curvature (trial ID p = .201), Stage 1 performance (trial ID p = .793), or Stage 2 performance (trial ID p = .531). When we considered the maximum escape performance values measured in each individual stickleback, we found that injection treatment was not correlated with a significant difference in either Stage 1 (p = 0.532) or Stage 2 (p = 0.102) performance (Figure 3). However, the mean Stage 2 performance among alum treated fish is higher, suggesting that there might be a trend if we account for other factors such as body stiffness and body curvature. Simply including these variables in a multivariate linear model did not improve our predictive power, as fibrosis severity score did not predict either Stage 1 performance (p = 0.509) or Stage 2 performance (p = 0.851). Instead, we turn to SEM to incorporate a hierarchical structure and further elucidate the effect of fibrosis on escape performance.

**Fig. 3.**
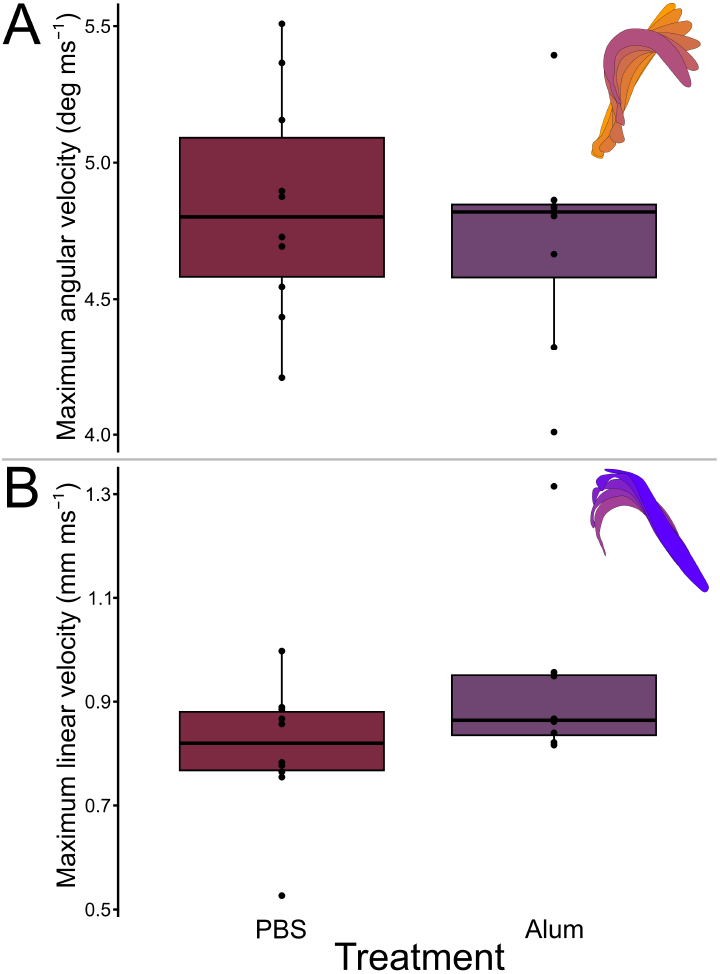
There were no significant differences between either Stage 1 (A) or Stage 2 (B) escape performance when compared between experimental treatments. However, fish treated with alum have a higher mean linear escape velocity during Stage 2 (P = 0.102, R2 = 0.158).

After running each of the hypothesized SEM models we were able to eliminate all but one based on the fit indices, since most models had several indices outside the threshold of acceptable values (Table 1). This left us with the model that included the base path as well as a behavioral effect connecting fibrosis and body curvature. This model selection was confirmed by both AIC and BIC (Table 1), and therefore all further results are obtained from this model. We find that all individual connections within this model are significant (p < 0.05) with the exception of the effect of body curvature on angular velocity (p = 0.656, Table 2; Figure 4G) and the effect of treatment on the interaction term (p = 0.503, Table 2, Figure 3). This means that fibrosis does not affect angular turning (Stage 1) performance through any of our hypothesized mechanisms.

**Table 2.** Model results of the Akaike’s information criterion (AIC) and Bayesian information criterion (BIC) selected path. Standardized coefficient estimates represent the increase in standard deviations of the response variable given an increase of one standard deviation in the explanatory variable. Coefficient estimates are in the units of the response variable over the explanatory variable. Correlations with a p-value >0.1 are listed in the table with gray text but are still part of the overall SEM model.

| Response Variable | Explanatory Variable | Standardized Coef. Estimate | Coef. Estimate | Std. Error | Z-Value | P-Value |
| --- | --- | --- | --- | --- | --- | --- |
| Fibrosis Score ($R^2=.92$) | | | | | | |
| | Treatment | 0.96 | 0.74 | 0.05 | 13.9 | $p<.001$ |
| Body Stiffness ($R^2=.23$) | | | | | | |
|  | Fibrosis Score | 0.48 | 0.36 | 0.16 | 2.34 | 0.02 |
| Treat. interaction ($R^2=.668$) | | | | | | |
|  | Treatment | -0.1 | -0.05 | 0.07 | -0.67 | 0.503 |
| | Stiffness | 0.86 | 0.65 | 0.12 | 5.62 | $p<.001$ |
| Body Curvature ($R^2=.50$) | | | | | | |
|  | Fibrosis Score | -0.63 | -0.75 | 0.23 | -3.29 | 0.001 |
| | Body Stiffness | 1.11 | 1.75 | 0.49 | 3.54 | $p<.001$ |
|  | Treatment Interaction effect | -0.9 | -1.89 | 0.61 | -3.1 | 0.002 |
| Stage 1 Perf. ($R^2=.01$) | | | | | | |
|  | Body Curvature | 0.104 | 0.09 | 0.2 | 0.45 | 0.656 |
| Stage 2 Perf. ($R^2=.29$) | | | | | | |
|  | Body Curvature | -0.54 | -0.17 | 0.064 | -2.72 | 0.007 |
| Total Effects |  |  |  |  |  |  |
| Stage 1 Perf. |  |  |  |  |  |  |
|  | Stiffness (PBS) | 0.12 | 0.16 | 0.35 | 0.44 | 0.658 |
|  | Fibrosis Score (Alum) | -0.05 | -0.05 | 0.11 | -0.44 | 0.661 |
| Stage 2 Perf. |  |  |  |  |  |  |
|  | Stiffness (PBS) | -0.6 | -0.3 | 0.141 | -2.16 | 0.031 |
|  | Fibrosis Score (Alum) | 0.253 | 0.1 | 0.05 | 1.81 | 0.071 |
| Indirect effects |  |  |  |  |  |  |
| Stage 2 Perf. |  |  |  |  |  |  |
|  | Fibrosis Score (Alum, via behavior) | 0.34 | 0.13 | 0.06 | 2.1 | 0.036 |
|  | Fibrosis Score (Alum, via stiffness) | -0.09 | -0.03 | 0.03 | -1.12 | 0.264 |

**Table 3.** Results of the Rosner test of multiple outliers (EnvStats V2.4.0 in R V4.0.2) reveal that four points are statistical outliers and were therefore removed from further analysis.

| Mean.i | SD.i | Value | Obs.Num | R.i+1 | lambda.i+1 | Outlier |
| --- | --- | --- | --- | --- | --- | --- |
| -3.57288E-17 | 0.5862627 | -1.500067 | 13 | 2.558694 | 2.858923 | TRUE |
| 0.05769487 | 0.513792 | -1.355515 | 2 | 2.75055 | 2.840774 | TRUE |
| 0.1142233 | 0.4340941 | 1.403849 | 14 | 2.970844 | 2.821681 | TRUE |
| 0.06048887 | 0.3482922 | 1.13918 | 5 | 3.097087 | 2.801551 | TRUE |

**Fig. 4.**
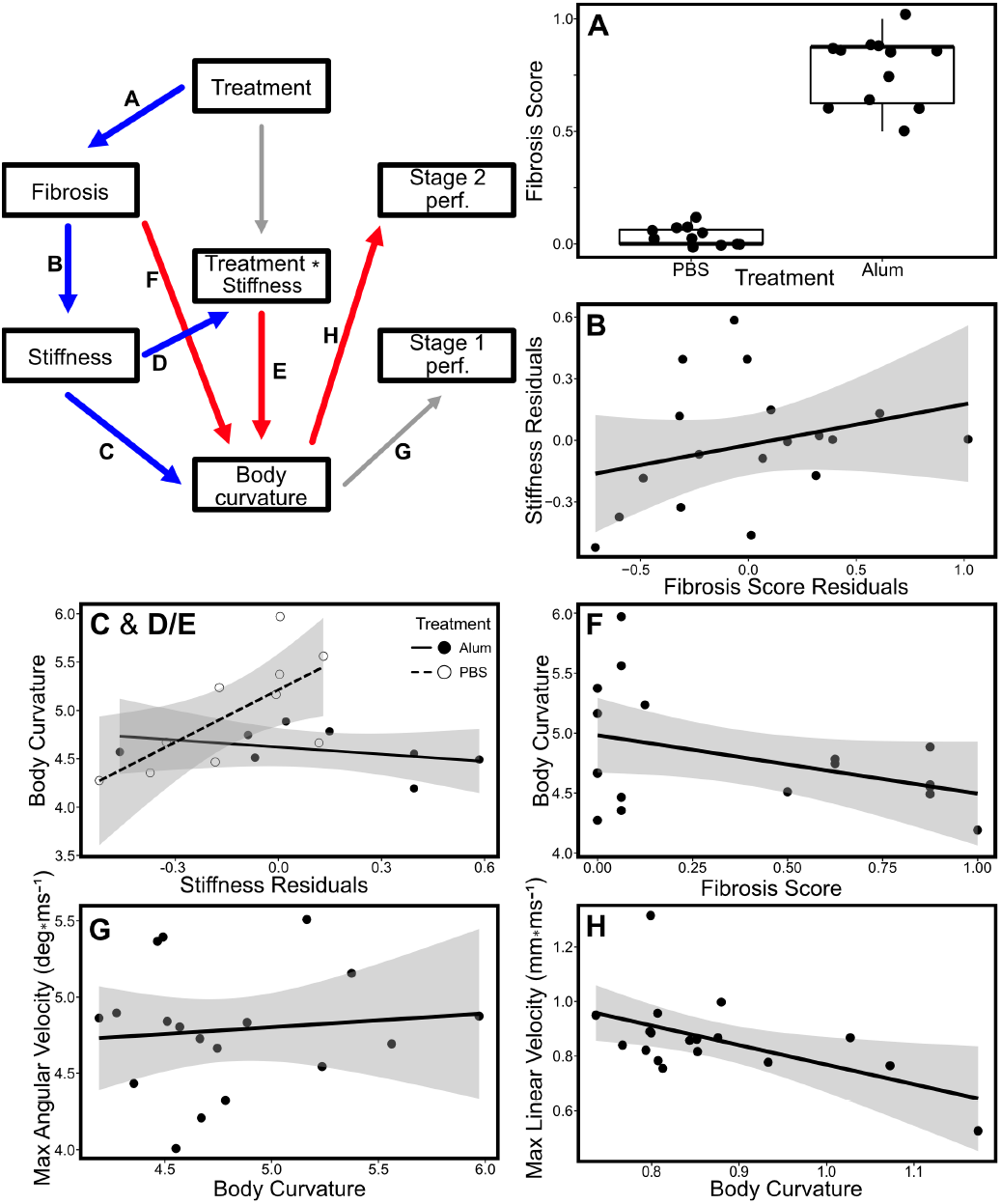
The best model to describe the effect of the fibrosis immune response on C-start escape performance included the base path as well as a behavioral effect and an interaction between treatment and stiffness. Regressions represent the individual paths linking Treatment, fibrosis severity score, body stiffness, body curvature, Stage 1 performance, and Stage 2 performance. The regression in B uses the residuals of F to account for the use of fibrosis as an independent variable in two regressions. The interaction effect wherein treatment mediates the relationship between stiffness and body curvature is represented by two separate regression lines in C and D/E. Blue lines represent positive coefficients, red lines represent negative coefficients, and gray lines represent non-significant correlations.

We find that fish injected with PBS have little to no fibrosis (Figure 4A), and therefore we do not present the effect of fibrosis on escape performance within these fish. However, we do find that these fish still show substantial variation in body stiffness and that the effect of stiffness on body curvature differs from the relationship seen in alum treated fish (Figure 4C, 4D/E)). Therefore, we present the effects of stiffness on escape performance within PBS fish to provide a point of comparison for the effects of fibrosis on performance. We find that stiffness is positively correlated with body curvature (p < 0.001, Table 2; Figure 4C, dashed line), which in turn is negatively correlated with Stage 2 performance (−0.17mm*ms-1, p = 0.007, Table 2; Figure 4H). Therefore, the total effect of stiffness on Stage 2 performance is a decrease of -0.6mm ms-1 for each standard deviation change in stiffness (p = 0.031, Table 2). Compared to the peak velocity in a PBS treated individual (0.998 mm ms-1), this equates to a 19.6% reduction in linear escape velocity from the least stiff PBS fish to the stiffest PBS fish. Unlike our control population of stickleback, fish injected with alum had high levels of fibrosis with greater variation between individuals (Figure 4A). Interestingly, while the high levels of fibrosis did increase body stiffness (p = 0.02, Table 2; Figure 4B), this was not correlated with a change in body curvature among alum treated fish (Figure 4D/E, solid line). Specifically, alum treated fish had relatively low body curvature regardless of their stiffness. This is further reflected in the fact that fibrosis levels are negatively correlated with body curvature (p = 0.001, Table 2; Figure 4F). Since body curvature is negatively correlated with Stage 2 performance (Table 2; Figure 4H), the maximum level of fibrosis corresponds with a 0.1 mm ms-1 increase in linear escape performance. This equates to a 12.3% increase in escape performance compared to the average fish treated with PBS. Since there was little variation body curvature due to stiffness among alum fish, we find that this performance increase is driven by a behavioral effect (p = 0.036, Table 2) instead of a change in mechanical properties (p = 0.264, Table 2).

## Discussion

### Performance effects

We initially asked what the effect of fibrosis was on C-start escape performance. Although classical methods of multivariate linear modeling were unable to uncover an effect, by using structural equation modeling (SEM) we were able to model this complicated causal relationship more completely. Using this method, we still found that there was no impact on the angular acceleration at the beginning of an escape response (Stage 1 performance) (Table 2). However, our best fit structural model indicated that natural variation in stiffness among fish without fibrosis negatively affects performance during the second stage of a C-start escape response (Table 2). Interestingly, this result did not apply to experimental fish that had fibrosis. Instead, we found that variation in stiffness among these fish had no effect on escape performance (Table 2), and that fibrosis affected linear escape performance by directly impacting body curvature independently of body stiffness. Since there was a negative correlation between both fibrosis and body curvature as well as body curvature and Stage 2 performance, we found that fibrosis acts to increase linear escape performance by as much as 12.3% in alum treated fish. Thus, in fish without fibrosis there is an inherent cost of having a stiff body in terms of linear escape performance, where stiffness can reduce Stage 2 velocity by as much as 20%. Interestingly, the presence of fibrosis and the concomitant high stiffness values appear to allow the fish to escape this cost and even improve escape performance. Past studies of stiffness and C-start performance showed that increases in stiffness tend to decrease Stage 1 performance and increase Stage 2 performance during a C-start escape response (Currier and Modarres-Sadeghi 2019). However, once a critical stiffness value is reached, Stage 2 performance decreases. We found that across both our experimental and control populations, Stage 1 performance variation did not align with these expectations since we did not find any correlation with body stiffness. Given that stickleback have naturally high body stiffness compared to other fishes (Witt et al. 2015), it is not surprising that during Stage 2 of the escape performance there would be a negative correlation between stiffness and escape performance. Indeed, this is exactly what we found in our control population. However, among fish treated with alum we find that there is no relationship between stiffness and Stage 2 escape performance. Surprisingly, we find that these fish have increased performance through another mechanism entirely. Although our data do not allow us to identify the alternate mechanism by which fish with fibrosis decrease body curvature thereby increasing Stage 2 escape performance, we suggest two possible causes. First, because our measurement of stiffness only accounted for passive body stiffness, it is possible that active changes to body stiffness were responsible for changes in body curvature that led to altered escape performance. Indeed, fish can increase their body stiffness by a factor of two through antagonistic muscle contraction (Tytell et al. 2018). Given that increased severity of fibrosis was correlated with increased passive body stiffness, it is possible that it also correlated with the strength of the lateral body musculature due to increased resistance to general movement. Currier and Modarres-Sadeghi (2019) suggested that active stiffness modulation could be an important mechanism through which fish overcome tradeoffs in escape performance, so this increased musculature could give fish with fibrosis a greater range of stiffnesses that can be achieved through behavioral modification. However, if the more flexible control fish were already reaching the critical stiffness at which Stage 2 performance decreases, then the ability to actively generate even higher stiffnesses might not be beneficial. Alternatively, the decreased body curvature in individuals with high fibrosis could be due to discomfort caused by the change in tissue composition of their coelom. Since the development and presence of fibrosis affects many mechanical as well as architectural properties of the body tissues (Wells 2013), it is possible that changes in tissue density or arrangement make high degrees of body curvature uncomfortable but don’t directly affect stiffness. Our results obtained from Stage 1 of the C-start are perhaps even more perplexing than those from Stage 2. Generally, we think of high body stiffnesses as having steep costs in Stage 1 performance, but the wide range of stiffnesses observed in our study did not seem to affect this part of the behavior at all. Although we considered an alternate model in which stiffness directly affected both performance metrics (Supplemental Figure 1C), all of our fit indices indicated that this model did not represent the data well (Table 1). Therefore, we think it is unlikely that Stage 1 performance is well explained by a different path among our measured variables. Additionally, active stiffness modulation is unlikely to explain this result because this only allows fish to actively increase body stiffness, but lower body stiffnesses tends to be associated with higher Stage 1 performance (Domenici 2001; Currier and Modarres-Sadeghi 2019). It is possible that stickleback are so stiff that Stage 1 behaviors have reached a performance trough, but unfortunately our data do not allow us to address this.

### Evolutionary implications

Given that stickleback from different freshwater populations have divergent immune responses with some expressing fibrosis and others suppressing it (Weber et al. 2022), the adaptive landscape of parasite defense is likely complex. We expected to find biomechanical costs of the immune response that would help explain why some populations suppress fibrosis. However, we did not identify any biomechanical costs in terms of escape performance and instead found a benefit. Importantly, we note that our study only examined C-start escape responses and fibrosis could carry biomechanical costs for other behaviors. For example, male courtship displays and nest-fanning behaviors entail rapid caudal vibration that could be impaired by the presence of fibrosis. However, given our results, we cannot conclude that biomechanical costs have any role in the suppression of fibrosis among these populations of stickleback. One of the primary functions of the C-start escape is to help fish evade predation. Interestingly, the cestode parasite completes its life cycle within bird predators of stickleback, and they are known to cause behavioral modifications in their hosts that leave the host more vulnerable to avian predation (Grécias et al. 2016, 2020; Berger and Aubin-Horth 2020). Although we do not suggest that fibrosis is under selection as a means of predator avoidance, we do find that it may help fish avoid predation in addition to suppressing cestode growth and viability. It is of particular note that fibrosis improves Stage 2 escape performance since this is the more salient part of the escape response in avoiding aerial predators. Importantly, this work represents the first demonstration of biomechanical locomotor effects of an immune response. The discordance of our result with past work on the relationship between material properties and escape performance suggests that the functional implications of immune responses may be more complicated than we initially predicted. Instead of simply considering the material property changes associated with fibrotic tissue, it may be important to consider other factors such as its effect on behavioral traits like active control of body stiffness. Additionally, it is likely important to consider the ecology of the fish and its parasite as this may provide important context for the desired functional outcomes of any immune response.

### Path analysis and structural equation modeling in evolutionary biology

Path analysis shares a long history with the field of evolutionary biology, having been initially applied by Sewall Wright (Wright 1918, 1921, 1934, 1960b,a). In the ensuing years this framework was expanded to include SEM, which uses unmeasured, inferred (latent) variables to model error terms (Reviewed in Tarka 2018). More recently, Stevan Arnold called for the use of path analysis as a means to connect phenotype to fitness with function as an intermediate (Arnold 1983). Despite this, relatively few studies have made use of these techniques within the fields of evolutionary and organismic biology (Matthews et al. 2023). We believe that this is an important technique that has great potential to advance these fields, and our study is just one example of how it can be used. Many functional morphology studies use classical linear regression modeling to show relationships between form and function. We initially conducted our data analysis in this way but were unable to find correlations between many of our variables. The only correlations that we did find were between variables that were expected to be highly related and would later end up adjacent in our SEM analysis. For instance, we could not answer our main question of whether fibrosis affects escape performance with traditional univariate or multivariate linear modeling. It was only by including a hierarchical structure with SEM analysis that we were able to pick out the broader effects of fibrosis on performance. By using some variables as intermediary steps, we were able to reduce noise in our estimate and find the true effect of fibrosis on the escape response. In addition to illuminating an effect that was obscured when using more common statistical methods, SEM allowed us to better understand the causation behind observed effects. For instance, we found that fibrosis acted to significantly increase Stage 2 escape performance in alum treated fish. However, it was only by calculating the indirect effects (Table 2) that we could show that this was due to a direct correlation between fibrosis and body curvature. We initially hypothesized that stiffness would be a critical intermediate that would facilitate this effect, however, the indirect effects revealed that this was not the case in our experimental population. Finally, although we did not employ it in our analysis, we believe that latent variable modeling is another noteworthy advantage of SEM. While latent variables are an inherent part of SEM for modeling error terms, they can also be used as independent variables in the model. This is particularly useful in organismal and evolutionary biology where many variables are known to be mechanistically important, but their measurement is not experimentally tractable. We suggest that future studies could use SEM with latent variables to include important links in their hypothesized path without needing to directly measure those traits. By combining the increased ability to model complex interactions granted by path analysis with the broader set of variables that can be included through latent variables, SEM can be an invaluable tool for biologists approaching any complex system.

## Acknowledgments

We thank Stephen DeLisle for help with injections and confirmation of fibrosis scores. We also thank Cory Hahn for his animal husbandry work. This research was supported by funding from Harvard University and an NSF graduate research fellowship under Grant No. DGE1745303 to DGM, Office of Naval Research Grants N00014-21-1-2210, N000141410533, and N00014-15-1-2234, NSF Grant 093088-17158 to GVL, and NIAID grant 1R01AI123659-01A1 to DIB. All procedures involving animals were reviewed and approved by the Institutional Animal Care and Use Committees of Harvard University and the University of Connecticut.

## Data availability statement

All data amd code used to produce these results will be made available upon final publication of the manuscript.

## Supplement

Supplemental video:

Example high speed video showing a ventral view of the dropped weight stimulus and an ensuing escape response.

https://www.youtube.com/watch?v=1xp_aalgLfs

**Supp. Fig. 1.**
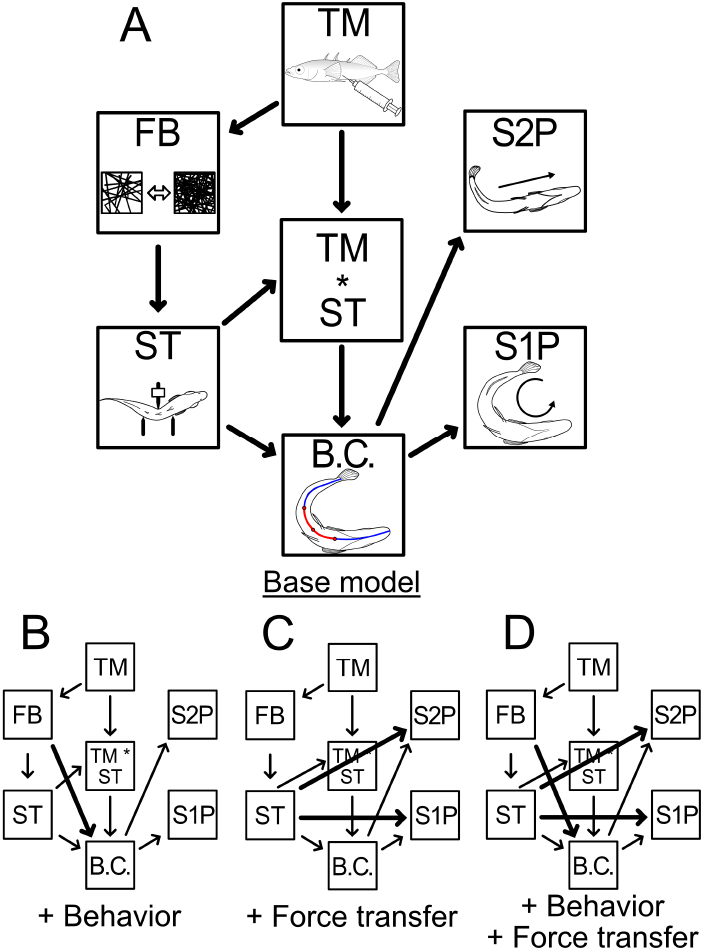
We test four alternative hypotheses of the causal relationship between fibrosis (FB) and both Stage 1 escape performance (S1P) and Stage 2 escape performance (S2P). We hypothesized that the level of fibrosis is determined primarily by the treatment (TM) that a fish receives and that the effect on performance is mediated by body stiffness (ST) and body curvature (B.C.) during an escape response. Additionally, we find that the relationship between stiffness and body curvature is mediated by an interaction with treatment. Bold lines represent causal relationship added in addition to the base model. These are thought to represent effects due to behavioral changes and due to the physical ability of the body to transfer force to the water.

**Supp. Fig. 2.**
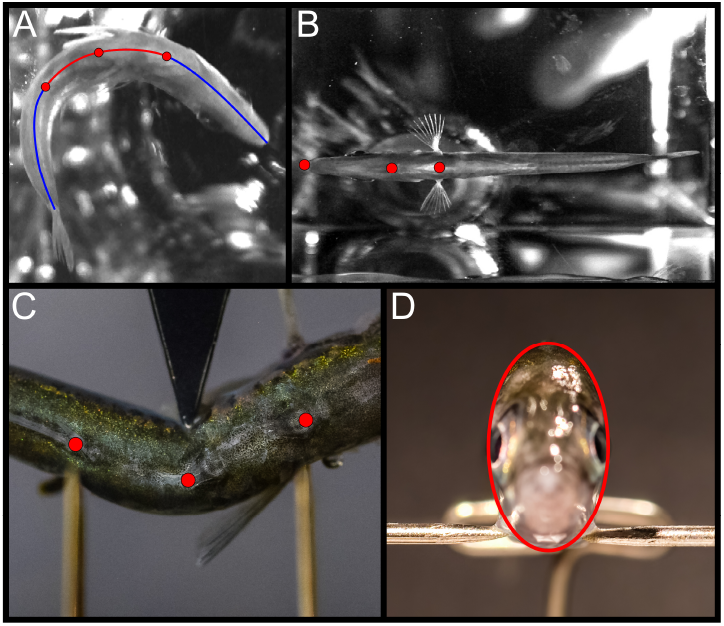
Example digitization locations (points) for all analyses. A) To measure body curvature during a C-start we traced full midlines from the snout to the caudal peduncle (blue line). These were then resampled to get the curve of the middle third (red line), specifically the points 1/3, 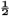, and 2/3 along the initial midline (red dots). B) We measured performance by digitizing points corresponding to the snout, posterior-most overlapping point of branchiostegal rays, and the midpoint between the pectoral fins. C) stiffness was measured by digitized points along the fish’s midline at the position corresponding to each of the three arms in our three-point bending test. D) We measured cross sectional area by taking head-on photos of euthanized fish and tracing an oval around the widest point of the body.

